# Supercharged Protein Nanosheets for Cell Expansion on Bioemulsions

**DOI:** 10.1101/2022.06.21.497058

**Authors:** Alexandra Chrysanthou, Hassan Kanso, Wencheng Zhong, Li Shang, Julien E. Gautrot

## Abstract

Cell culture at liquid-liquid interfaces, for example at the surface of oil microdroplets, is an attractive strategy to scale up adherent cell manufacturing whilst replacing the use of microplastics. Such process requires the adhesion of cells at interfaces stabilized and reinforced by protein nanosheets displaying high elasticity, but also presenting cell adhesive ligands able to bind integrin receptors. In this report, supercharged albumins are found to form strong elastic protein nanosheets and mediate extracellular matrix (ECM) protein adsorption and cell adhesion. The interfacial mechanical properties and elasticity of supercharged nanosheets is characterized by interfacial rheology and behaviors are compared to those of native bovine serum albumin, human serum albumin and α-lactalbumin. ECM protein adsorption to resulting supercharged nanosheets is then quantified via surface plasmon resonance and fluorescence microscopy, demonstrating the dual role supercharged albumins are proposed to play, as scaffold proteins structuring liquid-liquid interfaces and substrates for the capture of ECM molecules. Finally, the adhesion and proliferation of primary human epidermal stem cells is investigated, at pinned droplets, as well as on bioemulsions stabilized by corresponding supercharged nanosheets. This study demonstrates the potential of supercharged proteins for the engineering of biointerfaces for stem cell manufacturing, and draws structure-property relationships that will guide further engineering of associated systems.

## 1. Introduction

Tissue culture plastics, glass and other rigid substrates remain the main substrates on which adherent eukaryotic cells are routinely cultured and expanded. However, the use of such substrates constitutes an important hurdle to the scale up of cell manufacturing processes and alternative microplastics used for implementation in 3D bioreactors ^[1][2]^ pause increasing concerns in terms of contamination of cell products and to their processing. Liquid substrates, such as oil microdroplets, appear attractive alternatives for such 3D culture and manufacturing scale up ^[3]^. Indeed, it was recently proposed that the formation of mechanically strong protein or polymer nanosheets self-assembling at liquid-liquid interfaces and stabilizing oil microdroplets could provide a suitable local mechanical environment sustaining cell adhesion, whilst enabling readily processing of corresponding bioemulsions via centrifugation or potentially filtration ^[4][5]^. In particular, interfacial viscoelasticity of corresponding liquid-liquid systems was found to be regulating the ability of adherent stem cells to proliferate at corresponding liquid substrates ^[6]^. Although the ability of amphiphilic molecules and proteins, including globular proteins such as albumins, to stabilize liquid-liquid interfaces and act as tensioactive agents is well documented, the chemical and structural parameters regulating their interfacial viscoelasticity remains poorly understood.

Serum albumins are the most abundant proteins present in systemic circulation and play a significant role in the maintenance of osmotic pressure and pH of the blood ^[7][8]^. These albumins can bind a variety of substrates, including metal ions, fatty acids and therapeutic molecules, and have found broad applications in biotechnologies ^[9][7] [8]^. Owing to their hydrophobic core, serum albumins can bind water insoluble small negatively charged hydrophobic molecules which can regulate various interactions and functions such as the transportation of fatty acids in the blood ^[10]^. In addition, this inherent amphiphilicity has led to their wide application for the stabilization of emulsions and the formulation of food and healthcare products, and control of their rheological properties ^[11]^.

Human serum albumin (HSA) displays a comparable molecular weight to that of bovine serum albumin (BSA), near 66 kDa ^[12]^, and high percentage of homology (76%) ^[8]^. Both albumins display hydrophobic pockets allowing the binding of lipids and hydrophobic interfaces, and 17 disulfide bridges controlling the overall shape of these predominantly α-helical proteins. In contrast, α-lactalbumin (αLA) is significantly smaller, with a molecular weight of 14.2 kDa, and displays a relative abundance of lysine, cysteine, tyrosine and tryptophan residues ^[13] [14]^. α-lactalbumin participates in the binding of fatty acids or small molecules as for other albumins, and contributes to lactose synthesis ^[15][13]^. Yuan et al, (2018) reported enhanced antioxidant properties of the alpha-lactalbumin after ultrasound or enzymatic treatment possibly due to the formation of lactase crosslinked product with improved mechanical properties ^[15][16] [17]^. This suggested that modification of albumins may result in the control of their physico chemical and mechanical properties.

Chemical modifications of proteins can lead to significant structural changes and can help in understanding the role of electrostatic interactions on their stability. For example, modifications such as succinylation and acetylation can expose, or block, reactive amino acids in a protein, affecting protein-protein interactions, normal unfolding and aggregation^[18] [19] [20][21]^. The extent to which the structure of the protein is affected by such external factors is often determined by the natural rigidity of the protein. Flexible proteins such as caseins are relatively resistant to significant conformational changes in comparison to BSA and whey proteins which often experience high degrees of denaturation upon acylation. In contrast, chemical modifications may enhance the flexibility of globular proteins and offer potential for further conjugation ^[19]^. In turn, increasing charge density at the surface of proteins, using polyelectrolyte block copolymers for example, can enhance their adsorption at liquid-liquid interfaces, as in the case of chloroform-water interfaces ^[22]^. Similarly, aggregation at the surface of nanoparticles can be promoted by such modifications and associated changes in surface charges ^[23]^, modulating colloidal stability and physico-chemical properties. This implies a role for the charge density of proteins for the regulation of physicochemical and mechanical properties of resulting assemblies.

Supercharged proteins, naturally occurring, engineered or resulting from chemically modified native proteins, are attractive building blocks to design soft matter with emerging properties ^[24]^. Their engineering enabled the control of colloidal assembly ^[25]^, the formation of coacervates with RNA and other polyelectrolytes ^[26]^ and the formation of nanostructured films ^[27]^. The architecture of supercharged proteins ranges from well-structured and folded, as in the case of β-barrel proteins such as engineered GFP, to disordered macromolecules such as histones, involved in the formation of coacervates and the structuring of DNA, to elastin like proteins regulating the assembly of hydrogels and polyelectrolyte films with controlled mechanical properties. Hence supercharged proteins appear as attractive candidates for the stabilization and structuring of liquid-liquid interfaces and the engineering of interfacial mechanics, whilst conferring high surface charge densities to resulting interfaces for subsequent ECM protein adsorption.

In this report, the formation of supercharged protein nanosheets self-assembled at liquid-liquid interfaces is described. Supercharged BSA, generated via chemical modification, was assembled at the surface of the cytocompatible fluorinated oil Novec 7500, and the mechanical properties of resulting nanosheets was characterized via interfacial rheology and compared to that of other albumins. The impact of charge density, coupled to the formation of physical quadrupolar crosslinks on interfacial shear moduli and viscoelasticity is studied. In turn, the ability to adsorb extra-cellular matrix (ECM) proteins at the surface of supercharged nanosheets is characterized and the impact of the combined interfacial viscoelasticity and ECM adsorption on epidermal stem cell adhesion and proliferation is studied. Our results demonstrate the tuning of interfacial mechanics and ECM adsorption at liquid-liquid interfaces through supercharged nanosheet assembly and the potential of this platform for the culture of stem cells on bioemulsions.

## 2. Results

To explore structure-property relationships connecting albumin architecture, self-assembly and mechanics of resulting nanosheets at liquid-liquid interfaces, we first investigated the impact of the molecular structure of different albumins (bovine serum albumin, BSA; human serum albumin, HSA; α-lactalbumin, αLA) on the interfacial rheology of corresponding interfaces (Figure 1). In the absence of any co-surfactant, all three albumins adsorbed at corresponding interfaces rapidly (plateaus reached within 20 min), with comparable kinetics. At equilibrium, corresponding nanosheets displayed interfacial shear storage moduli in the range of 10^−2^-10^−1^ N/m, in agreement with literature reports for BSA ^[28][4][6][29][30]^(Figure 1D-E and Supplementary Figure S1A). The strongly hydrophobic character of Novec 7500, compared to other oils investigated in the literature (e.g. alkanes and hydroxy-alkanes) may account for the relatively high moduli observed. Hence, BSA as well as other globular proteins were found to form softer nanosheets at oil interfaces displaying more polar architectures ^[28]^. Compared to HSA and αLA, BSA formed stiffer nanosheets, displaying higher interfacial shear storage moduli, (Figure 1E and Supplementary Figure S1A). In addition, BSA and HSA nanosheets displayed higher elasticities, determined from interfacial stress relaxation experiments, with stress retentions of 33.9 ± 1.3 and 49.4 ± 6.1 %, respectively, compared to 21.8 ± 5.5 % for αLA (Figure 1F and Supplementary Figure S2A). This was in agreement with the higher loss modulus observed for αLA compared to BSA and HSA, and the stronger frequency dependence of the interfacial storage modulus measured for this protein (Supplementary Figure S1A). However, relaxation constants associated with stress dissipation at corresponding interfaces remained relatively comparable (Supplementary Figure S3A), implying similar dimensionalities for the protein networks assembled.

**Figure 1.**
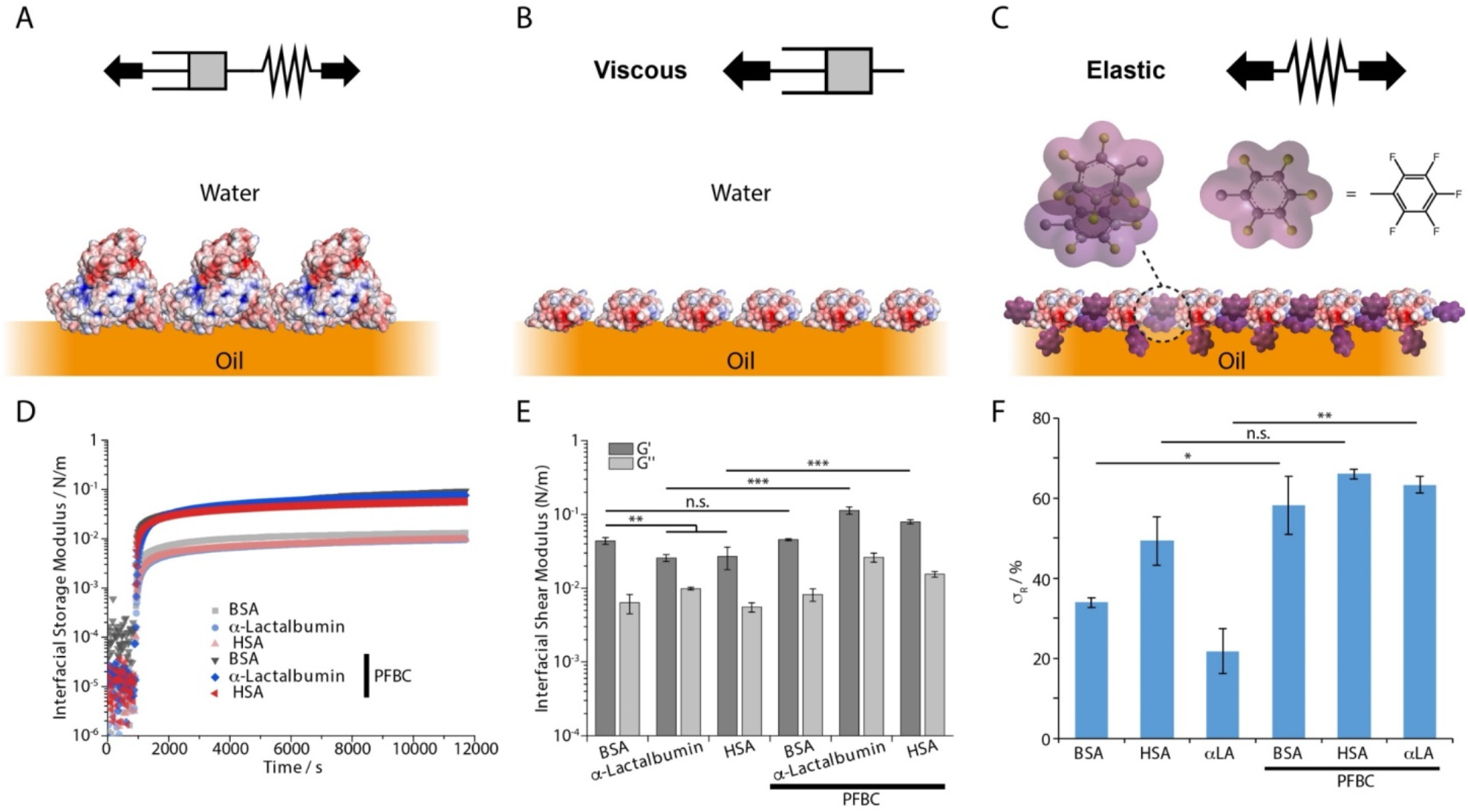
Formation and mechanical properties of albumin nanosheets at oil-water interfaces, in the presence of co-surfactant PFBC. (A) BSA/HSA form viscoelastic nanosheets at the oil-water interface. (B) α-Lactalbumin forms viscous nanosheets at the oil-water interface. (C) BSA, HSA and α-Lactalbumin form more elastic nanosheets in the presence of PFBC. (D) Time sweeps of the evolution of interfacial storage moduli of Novec 7500/PBS interfaces during the assembly of native BSA, HSA and α-Lactalbumin with and without co-surfactant PFBC (10 μg/mL in the oil phase). (E) Corresponding interfacial shear storage moduli extracted from frequency sweeps at oscillating amplitude of 10^−4^ rad. (F) Residual elasticities σ_R_ (%) extracted from the fits of stress relaxation experiments at 0.5 % strain.

The impact of the co-surfactant pentafluoro benzoyl chloride (PFBC) was explored next. PFBC had a significant impact on the viscoelastic properties of nanosheets generated from the three proteins studied (Figure 1 and Supplementary Figures S1-3). The interfacial storage moduli of nanosheets increased by almost one order of magnitude in the presence of PFBC (in the case of HSA and αLA, BSA displaying a more modest increase), as evidenced by frequency sweeps (Supplementary Figure S1 and Figure 1E). In addition, the elasticity of nanosheets formed in the presence of PFBC increased compared to nanosheets assembled in the absence of co-surfactant (Figure 1F and Supplementary Figure S2), presumably reflecting the impact that reactive co-surfactants such as PFBC play on physical crosslinking of nanosheets ^[6]^. This was also in agreement with the decrease in frequency dependency of the interfacial storage moduli of corresponding nanosheets, especially in the case of αLA (Supplementary Figure S1). Indeed, the increase in storage modulus and elastic stress retention σ_r_ in the presence of the co-surfactant PFBC was particularly pronounced in the case of αLA, switching from a relatively fluid, viscous interface, to a predominantly elastic interface (σ_r_ of 56.3 ± 2.1 %). The comparable mechanical properties of BSA and HSA nanosheets may be anticipated from the similarity of their molecular weight (66 and 64 kDa, respectively), amino acid composition (76 % homology) and isoelectric point (4.5 and 4.7, with zeta potentials of −20 and −21 mV, respectively; Supplementary Figure S4) ^[31][8][32] [33] [34][35]^. In contrast, αLA has a molecular weight of only 14 kDa and significantly different amino acid composition (36 % homology with BSA, only 31 % α-helix composition, Supplementary Figure S4) ^[36][37]^. Hence, we propose that the smaller size and more disordered structure of αLA results in more classic tensio-active properties, without the formation of protein networks at liquid-liquid interfaces (perhaps associated with reduced denaturation upon adsorption), resulting in more fluid interfaces with lower interfacial storage moduli, compared to BSA and HSA. However, in the presence of the co-surfactant PFBC, the abundance of functionalisable residues (e.g. lysine, serine, tyrosine, threonine) at the surface of the three types of albumins tested (see Supplementary Figure S4) underpinned the formation of physical crosslinks and the establishment of a more interconnected protein network, associated with an increase in interfacial elasticity (Figure 1F and Supplementary Figure S2-3).

Therefore, the amino acid composition and conformation of globular proteins not only regulate their assembly and interfacial mechanics at liquid-liquid interfaces, as was previously reported ^[38][6][29][30]^, but also impact on the response of these proteins to co-assembly with surfactants such as PFBC. These observations raise the possibility of engineering protein nanosheets via the design of their amino acid composition and chemistry. To further demonstrate this concept, we studied the impact of functionalization of BSA with succinic anhydride (leading to a negatively supercharged protein, aBSA, with a ζ-potential of −31.4 mV) and ethylene diamine residues (leading to a positively supercharged protein, cBSA, with a ζ-potential of +13.9 mV) ^[21]^ on self-assembly and interfacial mechanics (Figure 2). In the absence of co-surfactant, the surface chemistry of BSA had a striking impact on the mechanical properties of corresponding nanosheets assembled at Novec 7500-water interfaces. Supercharged proteins resulted in softer nanosheets, presumably as a result of increasing repulsion between proteins assembled at corresponding interfaces (Figure 2D-E and Supplementary Figure S1B). This effect was particularly pronounced in the case of aBSA that had a higher charge density compared to cBSA. Indeed, the interfacial storage modulus increased only weakly upon exposure to aBSA and was comparable to the interfacial loss modulus extracted from measurements. At equilibrium, the residual elastic stress measured from stress relaxation experiments was particularly low in the case of aBSA (Figure 2F and Supplementary Figures S2B) and relaxation profiles were associated with reduced rate constants (Supplementary Figure S3B).

**Figure 2.**
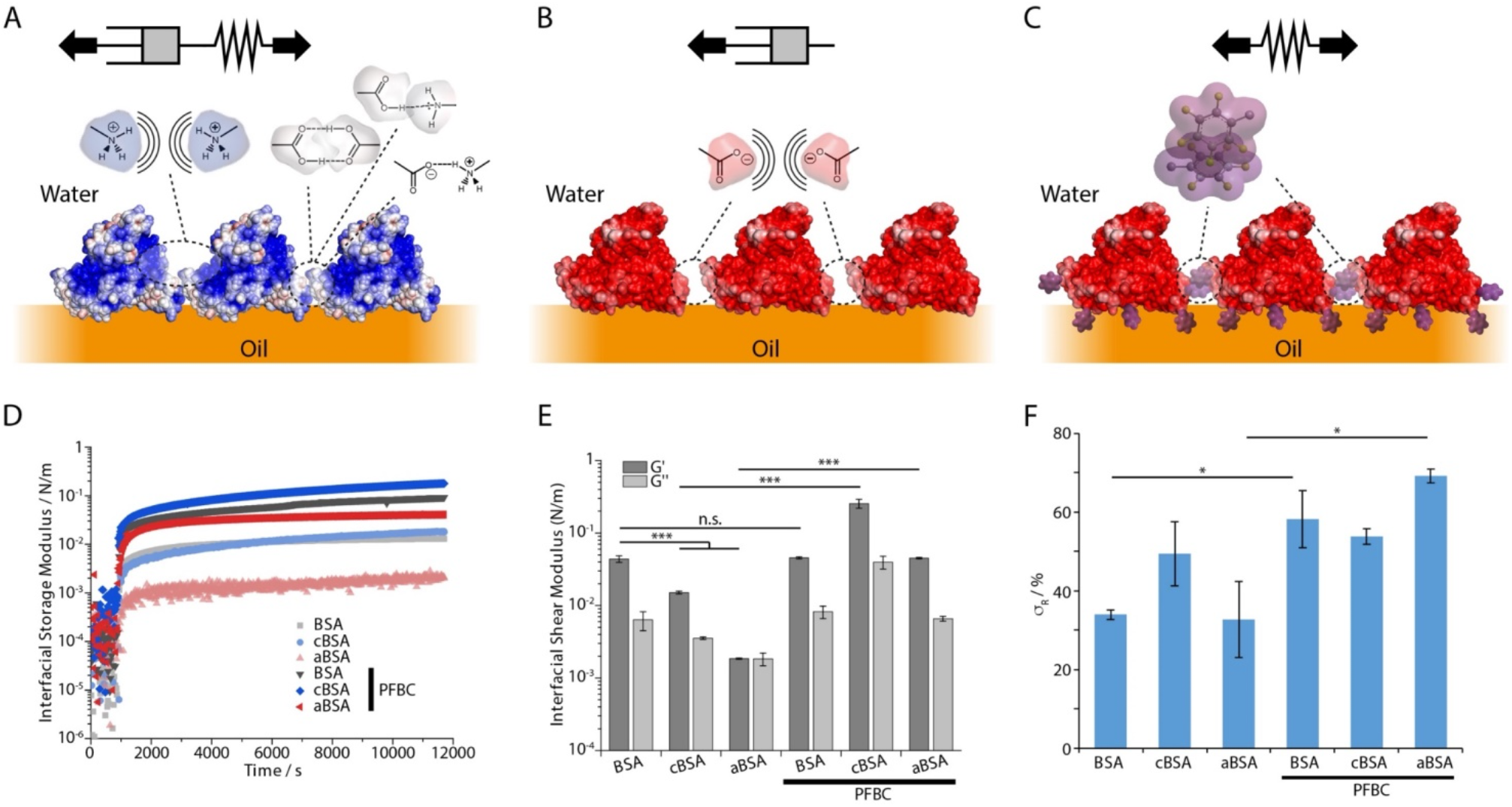
Formation and mechanical properties of supercharged albumin nanosheets at oil-water interfaces, in the presence of co-surfactant PFBC. (A) Cationic BSA (cBSA) forms viscoelastic nanosheets at the oil-water interface. (B) Anionic BSA (aBSA) forms viscous nanosheets at the oil-water interface. (C) In the presence of PFBC, supercharged albumins, including aBSA, form elastic nanosheets. (D) Time sweeps of the evolution of interfacial storage moduli of Novec 7500/PBS interfaces during the assembly of native BSA, cBSA and aBSA, with and without co-surfactant PFBC (10 μg/mL in the oil phase). (E) Corresponding interfacial shear storage moduli extracted from frequency sweeps at oscillating amplitude of 10^−4^ rad. (F) Residual elasticities σ_R_ (%) extracted from the fits of stress relaxation experiments at 0.5 % strain.

In contrast, cBSA formed slightly softer, but more elastic interfaces (Figure 2E-F and Supplementary Figure S2B). We confirmed that protein densities adsorbed to fluorophilic model interfaces (monolayers of perfluorinated alkyl thiols) were comparable between BSA and aBSA, and slightly higher for cBSA (Figure 3A). Therefore the relatively strong surface potential associated with supercharged BSAs is proposed to result in enhanced repulsion between albumin molecules assembling at liquid-liquid interfaces, without substantial reduction in surface densities, resulting in softer interfaces, compared to native BSA nanosheets, in particular in the case of aBSA (Figure 2A-C). However, the occurrence of hydrogen bonding, and potentially solution aggregation, in the case of cBSA led to an increase in elasticity of corresponding interfacial networks. The interfacial storage moduli measured were in between those of BSA and aBSA nanosheets (Figure 2E) and, after an initial rapid increase, a gradual increase in modulus was observed. This may reflect that the surface charge density of cBSA, although more modest than that of aBSA, results in initial repulsion between surface adsorbed macromolecules, but that with time further infiltration and interactions (perhaps between amine and carboxylic residues) result in physical crosslinking of associated nanosheets. The increased surface density of cBSA adsorbed at fluorophilic monolayers compared to BSA (Figure 3A), together with the increased elasticity of these interfaces, may also reflect the adsorption of small protein aggregates, rather than isolated proteins, that develop further interactions following adsorption. In agreement with such hypothesis, the frequency dependency of the interfacial storage modulus of cBSA nanosheets was moderate compared to that of aBSA nanosheets (Supplementary Figure S1B).

**Figure 3.**
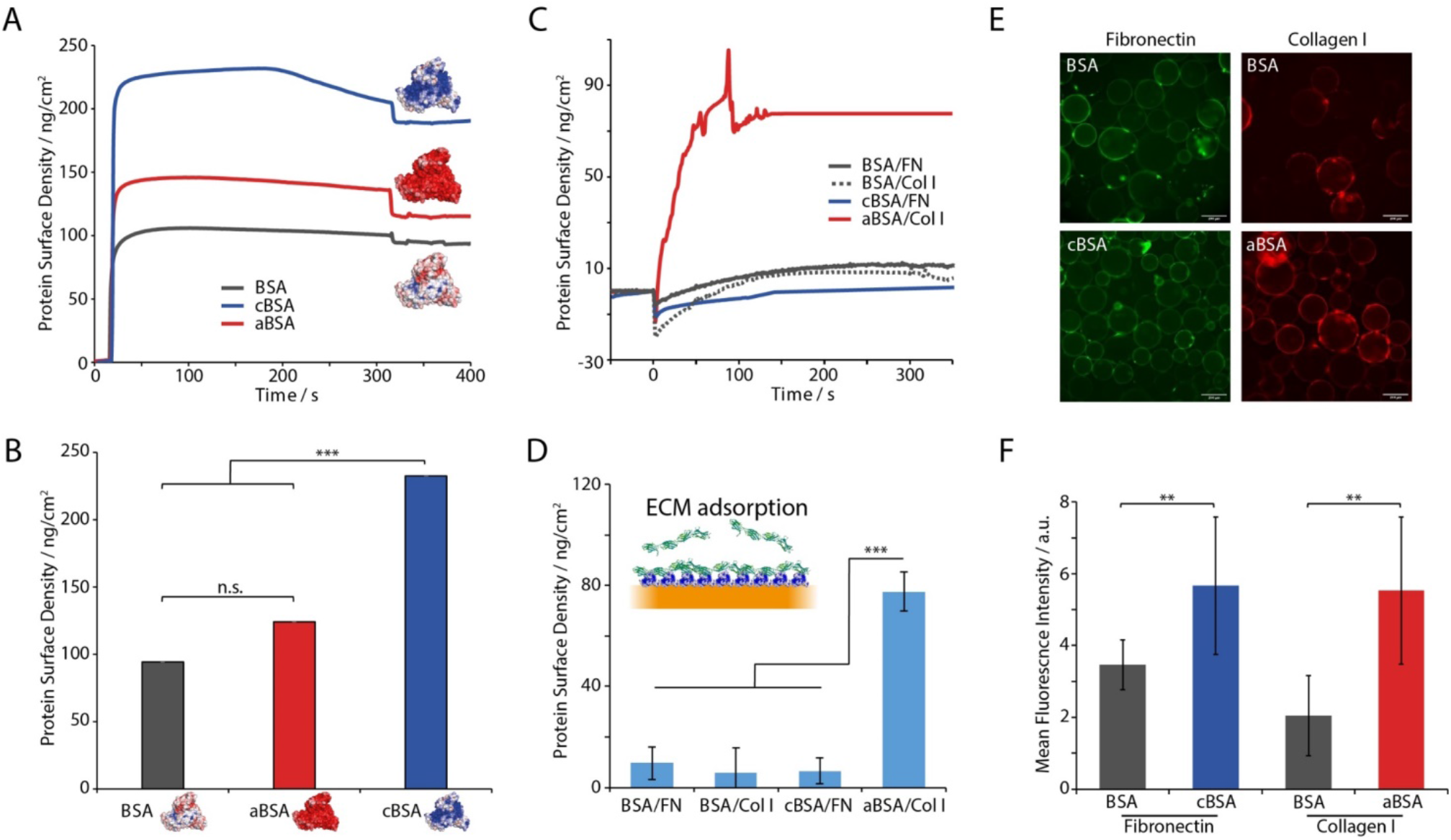
(A) Representative surface plasmon resonance traces of the adsorption of supercharged albumins and native BSA to perflurodecanethiol monolayers modelling fluorinated oil interfaces. (B) Corresponding quantification of resulting protein surface densities. (C) SPR quantification of fibronectin (FN) or collagen type I (Col I) adsorption at the surface of supercharged protein layers. (D) Corresponding protein surface densities. (E) Epifluorescence microscopy images of FN or Col I adsorption onto BSA, cBSA and aBSA emulsions; Green, FN; red, Col I. Scale bars, 200 µm. (F) Quantification (mean fluorescence intensity) of adsorbed FN or Col I on corresponding protein nanosheets. Error bars are s.e.m.; n = 3.

In the presence of the co-surfactant PFBC, supercharged proteins assembled into significantly stiffer nanosheets (Figure 2D-E). In particular, the interfacial storage modulus of cBSA and aBSA nanosheets was 17 and 24 fold higher, respectively, in the presence of PFBC, compared to only 5% increase for BSA. As for other albumins studied, the increase in hydrophobicity and associated physical crosslinks enabled by PFBC moieties resulted in stiffening of nanosheets. In this respect, the higher storage moduli measured for cBSA nanosheets may reflect a combined impact of hydrophobic crosslinks from strong quadrupole interactions between perfluorobenzene moieties ^[39]^ and electrostatic crosslinking or hydrogen bonding associated with interactions between amines and carboxylic moieties of cBSA macromolecules. In contrast, the high surface charge of aBSA prevented the formation of as extensive hydrophobic crosslinked networks compared to BSA nanosheets.

These changes in interfacial modulus are also supported by the increase of the elasticity (quantified through σ_R_) of all protein nanosheets in the presence of PFBC (Figure 2F), together with the increased relaxation times associated (Supplementary Figure S3B), in particular in the case of supercharged nanosheets. In this respect, the particularly striking shift of aBSA nanosheets towards a highly elastic network behavior, despite a weaker interfacial shear modulus compared to cBSA (in the presence of PFBC) suggests that its higher surface charge density and associated electrostatic repulsive forces may contribute to its stress relaxation. The increased elasticity evidenced in supercharged protein nanosheets formed in the presence of PFBC was also reflected in the reduction of the frequency dependency of their interfacial storage moduli (note the lack of decrease at high frequencies, see Supplementary Figure S1B).

Overall, supercharged protein nanosheets stabilised by the co-surfactant PFBC display a combination of strong interfacial mechanical properties and high surface potential, ideal to promote the adsorption of ECM proteins such as fibronectin and collagen, to enable cell adhesion. To quantify ECM protein adsorption to supercharged protein nanosheets, we first studied the formation of protein assemblies at the surface of model perfluorinated monolayers, via surface plasmon resonance (SPR, Figure 3A-B). Following protein injection, their adsorption rapidly increased, to reach a plateau near 90, 110 and 230 ng/cm^2^ for BSA, aBSA and cBSA, respectively. Such rapid adsorption is in line with the adsorption reported for albumin at other hydrophobic interfaces and gold substrates, and in agreement with the rapid increase in interfacial moduli observed via interfacial rheology ^[40][41][42][43]^. The higher rate at which protein adsorption was observed via SPR may be reflecting the fact that kinetics of evolution of interfacial mechanics not only depend on assembly at corresponding interfaces, but also the formation of physical crosslinks (via denaturation, hydrophobic bond formation, or electrostatic/hydrogen bonding) and a macroscale protein network. However, it is also likely that protein diffusion in the interfacial rheology trough is limiting protein adsorption to liquid interfaces, as was reported in other systems^[44]^.

Having examined the adsorption of supercharged albumins, we next quantified the secondary adsorption of ECM proteins to the resulting interfaces (Figure 3C-D). Fibronectin and collagen type I solutions (in PBS) were injected on cBSA and aBSA interfaces (respectively), on the basis of their low and high isoelectric points (6.0 and 8-9, respectively) ^[1][45][46]^. Adsorption levels were compared to those measured to native BSA interfaces. Collagen I adsorption was found to be enhanced on aBSA interfaces, supporting the hypothesis that enhanced charge density and associated electrostatic interactions, compared to native BSA, promotes the adsorption of high pI proteins. In contrast, fibronectin adsorption was moderate on both native BSA and cBSA, presumably as a result of the negative ζ-potential associated with native BSA, and the modest positive ζ-potential of cBSA compared to the negative potential achieved for for aBSA.

To quantify protein adsorption at the surface of supercharged nanosheets assembled at liquid-liquid interfaces, we generated nanosheet-stabilised microdroplets and quantified fibronectin and collagen I adsorption via immunostaining and fluorescence microscopy (Figure 3E-F and Supplementary Figure S5). Images of resulting droplets clearly demonstrated ECM protein adsorption at the surface of nanosheets and the enhancement of such adsorption at the surface of supercharged protein nanosheets, in agreement with SPR data. In particular, collagen I was again found to adsorb strongly at the surface of aBSA, compared to native BSA, but fibronectin adsorption was also found to be slightly promoted, compared to native BSA. Therefore, our data demonstrate that supercharged albumins not only allow the assembly of stiff, strong protein nanosheets at liquid-liquid interfaces, but also enable the adsorption of ECM proteins relevant to regulate cell adhesion and stem cell expansion.

The adhesion and spreading of human primary keratinocytes is typically mediated by integrins and regulate their fate decision ^[47][48]^. Collagen (in particular type I and IV) is typically used to promote keratinocyte adhesion and selection, although fibronectin is also often used, despite the lack of expression of associated integrin heterodimers in normal human interfollicular epidermis ^[49] [50] [51] [52]^. To explore the ability to use microdroplets as microcarriers for the expansion of adherent stem cells, we first generated pinned droplets, stabilised by supercharged albumins, followed by adsorption of complementary ECM proteins (FN, fibronectin for cBSA and Col I, collagen I for aBSA). Human primary keratinocytes were then cultured on the resulting droplets, enabling the quantification of cell densities at different time points (Figure 4 and Supplementary Figures S6-7). As a comparison, we seeded cells on tissue culture polystyrene (TCP) and on poly(L-lysine) (PLL) stabilized pinned droplets ^[5][6]^. After 3 days of culture, cell densities on PFBC/cBSA/FN stabilised interfaces were comparable to those measured on TCP and PFBC/PLL/FN controls. In comparison, densities measured on supercharged nanosheets formed in the absence of PFBC, or on native BSA nanosheets were significantly reduced compared to controls. Densities measured on PFBC/aBSA/COL were intermediate between those measured for native BSA nanosheets and the controls, despite the strong adhesion keratinocytes typically display for collagen I-coated interfaces. After culture for 7 days, increased cell densities were observed for all conditions. Importantly, all PFBC reinforced nanosheets (displaying strong interfacial mechanical properties) enabled comparable cell densities to the controls to be achieved. In contrast, supercharged nanosheets generated in the absence of PFBC and displaying softer, more viscous behaviors were clearly unable to sustain rapid keratinocyte expansion (Figure 4A/B). This effect was particularly marked in the case of native nanosheets, in agreement with the combined weakness of corresponding interfaces and their lack of ability to mediate collagen I adsorption.

**Figure 4.**
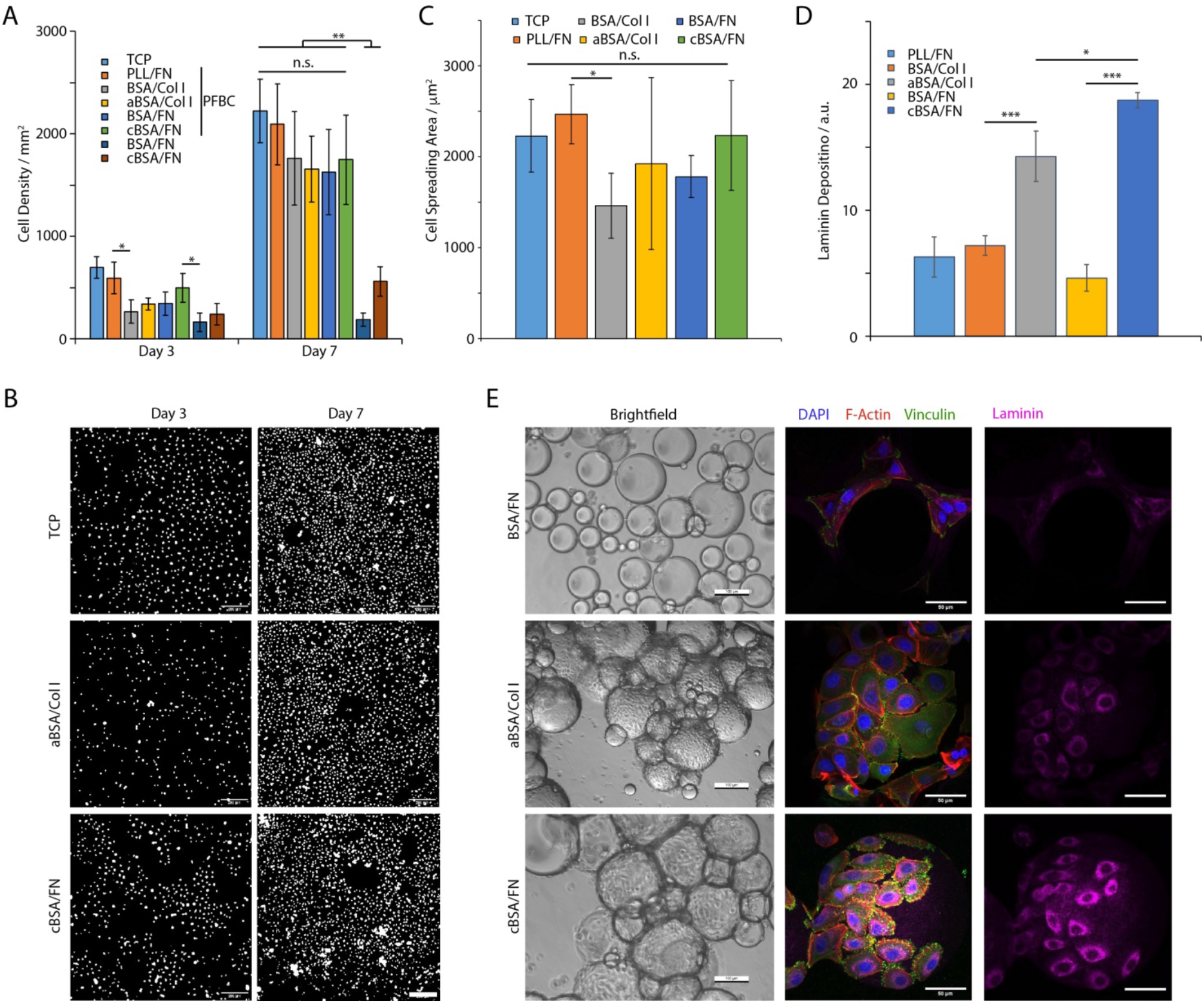
(A) Human primary keratinocytes (HPK) proliferation on interfaces conditioned with different supercharged albumins, functionalized with ECM proteins and assembled with or without co-surfactant PFBC. (B) Selected images of cells spreading at corresponding liquid-liquid interfaces after three and seven days of culture at TCP, cBSA/FN, aBSA/Col I with PFBC. Images are corresponding nuclear stainings. Scale bars are 200 µm. (C) Quantification of HPKs spreading area (24 h after seeding) characterized on pinned droplets functionalized with corresponding nanosheets. (D) Quantification of laminin deposition at liquid-liquid interfaces. (E) Bright field and confocal images of HPKs cultured for seven days on emulsions stabilised by protein nanosheets (blue, DAPI; red, phalloidin; green, vinculin; purple, laminin). Scale bars are 100 µm (bright field) and 50 µm (confocal). Error bars are s.e.m.; n = 3.

In agreement with these observations, cell spreading on supercharged nanosheets, reinforced by PFBC and with ECM protein adsorption, was comparable to controls, whereas those spreading on nanosheets generated from native BSA displayed more rounded morphologies, 24 h after seeding (Figure 4C). Therefore, our data indicate that cell adhesion to liquid interfaces reinforced by supercharged protein nanosheets displaying strong mechanical properties and high ECM adsorption enable rapid keratinocyte adhesion, spreading and expansion, comparable to what is typically observed on tissue culture plastic or at cationic polymer nanosheets (based on PLL). Over prolonged culture times, only reinforced nanosheets promoted keratinocyte expansion on pinned droplets.

The ability to expand keratinocytes at the surface of bioemulsions stabilized by supercharged protein nanosheets was examined next (Figure 4D/E). Bioemulsions were generated by enabling native BSA and supercharged nanosheets to stabilize microdroplets, prior to the assembly of corresponding ECM proteins. Interfaces not reinforced by co-surfactants did not sustain keratinocyte proliferation (Figure 4E and Supplementary Figures S8-9), in agreement with results obtained on pinned droplets. Too few cells were observed on these systems to enable further characterization. Few keratinocytes could be observed at the surface of droplets stabilized by native BSA, reinforced with PFBC, regardless of the ECM protein adsorbed (Figure 4E and Supplementary Figures S8-9). Similarly, keratinocytes adhering to supercharged protein nanosheets, in the presence of PFBC, displayed mature focal adhesions, assembled a well-structured actin cytoskeleton and deposited higher levels of laminin α1 compared to cell spreading at the surface of native BSA-stabilized droplets (Figure 4C/E and Supplementary Figure S10). Therefore, the more challenging adhesive environment associated with droplet curvature was met by the combination of high ECM protein adsorption and increased interfacial elasticity associated with supercharged protein nanosheets.

## 3. Discussion and Perspective

Focal adhesion formation is typically regulated by the rigidity of the substrate on which cells are adhering ^[53] [54][48][55]^. However, an increasing number of reports are suggesting that nanoscale mechanics, rather than bulk mechanical properties regulate such processes ^[56][57][48]^. This concept is taken to its extreme in the phenomenon of cell adhesion to liquid substrates such as fluorophilic or silicone oils, providing strong elastic polymer or protein nanosheets can be assembled at corresponding interfaces ^[4][5][58][6]^. Such phenomenon had remained restricted to a relatively small number of proteins and polymers and the translation of such systems to the culture of adherent cells on bioemulsions will require the engineering of more readily available macromolecules. Supercharged albumins appear as promising candidates for such applications, given their availability and amenability to engineering of chemical structure. Our data indicate that the combination of high interfacial modulus, elasticity and ability to promote ECM adsorption of surfactant-reinforced supercharged protein nanosheets enhances cell attachment and focal adhesion formation, despite the absence of underlying rigid or elastic substrate.

The engineering of scaffold proteins that display suitable combination of amphiphilicity, in order to adsorb at the surface and stabilize oil droplets, and interfacial mechanics, to sustain cell-mediated contractile forces, remains challenging. Despite the wealth of data describing the stabilization of emulsions by a range of proteins and surfactants, in particular for application in the food industry or the formulation of healthcare products, the emphasis is most often placed on surface tension and dilatational mechanics. Parameters impacting interfacial viscoelasticity, independent on changes in surface areas remain poorly understood and structure-property relationship enabling systematic design required. In this context, supercharged albumins were found not only to retain tensioactive properties suitable to stabilize microdroplets, but also retained the ability to couple with reactive co-surfactants such as PFBC in order to provide physical crosslinks required to achieve strong interfacial elastic properties. Therefore supercharged albumin nanosheets offer a suitable combination of interfacial scaffolding properties and ability to promote fast ECM protein adsorption without the need for further coupling or reactivity.

The application of such protein engineering to the design of bioemulsions suitable for adherent cell culture remains in its infancy. A range of stem cells have been cultured at such interfaces and the demonstration of long term expansion of mesenchymal stem cells on such expansion paved the way towards the translation of these systems to bioreactors ^[3]^. A broad range of parameters remains to be investigated, such as the ability to sustain matrix remodeling or how the control of droplet size, stability and curvature (which may affect cell adhesion, proliferation and fate decision ^[59]^) may also affect the quality of cells manufactured on resulting bioemulsions. However these systems offer a unique opportunity to replace plastics and microplastics and revolutionize cell manufacturing processes.

## 4. Materials and Methods

### Materials and Chemicals

Native, anionic and cationic BSA were prepared and provided as described in ^[21]^. The fluorinated surfactant 2,3,4,5,6,-Perfluorobenzoyl chloride, PBS, Trichloro (1H,1H,2H,2H-perfluorooctyl) silane (97%) and the 1H,1H,2H,2H-Perfluorodecanethiol (97%) were purchased from Sigma Aldrich Co. The fluorinated oil (Novec 7500) is from ACOTA. The SPR-Au chips were obtained from Ssens.

### Preparation of emulsions

Emulsions were generated using 1 mL of fluorinated oil (Novec 7500, ACOTA) with or without fluorinated surfactant (2,3,4,5,6-pentafluorobenzoyl chloride, PFBC, final concentration of 0.01 mg/mL) and 2 mL of protein aqueous solution (1 mg/mL in PBS), added to a glass vial. The vial was shaken for 15 s and incubate for 1 h at room temperature. The upper liquid phase (aqueous) was aspirated and replaced with PBS 6 times.

### Interfacial shear rheology measurements

Interfacial rheological measurements were carried out on a Discovery Hydrid-Rheometer (DHR-3) from TA Instruments, using a Du Nouy Ring geometry and a Delrin trough with a circular channel. The Du Nouy ring has a diamond-shaped cross section that improves positioning at the interface between two liquids to measure interfacial rheological properties whilst minimizing viscous drag from upper and sub-phases. The ring has a radius of 10 mm and is made of a platinum-iridium wire of 400 µm thickness. The Derlin trough was filled with 4 mL of fluorinated oil (with or without surfactant). Using axial force monitoring, the ring was positioned at the surface of the fluorinated oil, and was then lowered by a further 200 µm to position the medial plane of the ring at the fluorinated phase interface. 4 mL of the PBS solution were then gently introduced to fully cover the fluorinated sub-phase. Time sweeps were performed at a frequency of 0.1 Hz and temperature of 25°C, with a displacement of 1.0 10^−3^ rad (strain of 1 %) to follow the self-assembly of the protein nanosheets at corresponding interfaces. In each case, the protein solution (1 mg/mL) was added after 15 min of incubation and continuous acquisition of interfacial rheology data for the naked liquid-liquid interface. Before and after each time sweep, a frequency sweep (with displacements of 1.0 10^−3^ rad) and amplitude sweeps (at a frequency of 0.1 Hz) were carried out to examine the frequency-dependent behavior of corresponding interfaces and to ensure that the selected displacement and frequency selected were within the linear viscoelastic region.

### Surface plasmon resonance (SPR)

SPR measurements were carried out on a BIACORE X from Biacore AB. SPR chips (SPR-Au 10 × 12 mm, Ssens) were plasma oxidized for five minutes and then incubated in a 5 mM ethanolic solution of 1H,1H,2H,2H-Perfluorodecanethiol, overnight at room temperature. This created a model fluorinated monolayer mimicking the fluoriphilic properties of Novec 7500. The chips were washed once with water, dried in an air stream and kept dry at room temperature prior to mounting (within a few minutes). Thereafter, the sensor chip was mounted on a plastic support frame and placed in a Biacore protective cassette. The maintenance sensor chip cassette was first placed into the sensor chip port and docked onto the Integrated μ-Fluidic Cartridge (IFC) flow block, prior to priming the system with ethanol. The sample sensor chip cassette was then docked and primed once with PBS. Once the sensor chip primed, the signal was allowed to stabilize to a stable baseline, and the protein solution (1 mg/ mL in PBS) was loaded into the IFC sample loop with a micropipette (volume of 50 μL). The sample and buffer flow rates were kept at 10 μL/min throughout. After the injection finished, washing of the surface was carried out in running buffer (PBS) for 10 min. Washing of the surface was allowed to continue for 10 min prior to injection of the second protein (collagen or fibronectin at 10 μg/mL and 100 μg/mL in PBS, respectively; volume of 50 μL), at a flow rate of 10 μL/min. Buffer (PBS) was flown on the sensor chip for 10 min to wash off excess protein solution and data was allowed to continue for a further 10 min.

### Generation of fluorinated pinned droplets for cell culture

Thin glass slides (25 × 60 mm, VWR) were washed with isopropanol and dried under nitrogen, prior to plasma oxidation for 10 min (Henniker Plasma HPT-200; air). Slides were then placed into an anhydrous ethanol solution (9.5 mL) containing trichloro-1H,1H,2H,2H-perfluorooctyl silane (97%, Sigma) (500 μL) for 1 h, at room temperature. The fluorinated glass slides were cut into chips (1 × 1 cm) and placed into a 24 well plate (for Hoechst staining), or the glass slides were kept at their original dimensions and embedded on sticky-slide 8 wells plates (Ibidi), for imaging on a confocal microscope. After sterilization with 70 % ethanol, the wells were washed (once) and then filled with 2 mL (or 600 μL for the sticky wells) of PBS (pH 7.4 for the different BSA types and pH 10.5 for poly(L-lysine), PLL). 100 μL of fluorinated oil (or 10 μL for the sticky wells), with or without fluorinated surfactant (10 μg/mL) were added to the surface of the glass slide, forming a fluorinated pinned droplet. For samples prepared in 24 well plates, 30 μL of the oil phase was removed using a micropipette, to form a flatter oil droplet. For protein deposition, 10 μL of BSA solution (100 mg/mL) were added into PBS phase contained in the well (final concentration of 1 mg/mL; volume used for Ibidi well was only 8 μL) and incubated for 1 h. After the incubation time, wells were washed six times with PBS (by dilution/aspiration, ensuring the oil surface did not become exposed to air). Fibronectin (10 μg/mL, final concentration) or collagen type I (100 µg/mL, final concentration) were added into the PBS solution and incubated for 1 h, followed by washing with PBS (four times) and with keratinocyte basal medium 2 (KBM2, twice).

### Hoechst staining

Cell proliferation was assessed via Hoechst staining, microscopy and counting of nuclei. Cells were incubated in KBM2 containing 2 μL Hoechst 33342 (5 mg/mL stock solution, ThermoFischer Scientific) for 30 min before imaging by epifluorescence microscopy (see details below). The number of nuclei per image was determined manually, and converted in cell densities per surface area.

### Immuno-fluorescence staining and antibodies

Samples (emulsions) were washed (dilution and aspiration, followed by addition of solutions) once with PBS and fixed with 4 % paraformaldehyde (Sigma-Aldrich; 8 % for samples in Ibidi well plates) for 10 min at room temperature. Thereafter, samples were washed three times with PBS and permeabilized with 0.2 % Triton X-100 (Sigma-Aldrich; 0.4 % for samples in Ibidi well plates) for 5 min at room temperature. After washing with PBS (three times), samples were blocked for 1 h in 10 % fetal bovine serum. The blocking buffer was partly removed from the samples, not allowing them to be exposed to air, and the samples were incubated with primary antibodies at 4 °C overnight. Samples were washed six times with PBS and incubated for 1 h with the secondary antibodies (phalloidin, 1:500; DAPI, 1:1000; vinculin, 1:1000; laminin, 1:500) in blocking buffer (FBS 10% in PBS). After washing with PBS (six times), samples were transferred to Ibidi wells for imaging.

### Immuno-fluorescence microscopy and data analysis

Fluorescence microscopy images were acquired with a Leica DMi8 fluorescence microscopy. To determine cell densities per mm^2^, cell counting was carried out by thresholding and watershedding nuclei images in Fiji ImageJ. In the case of cell clusters, for which this method did not allow the isolation of individual nuclei, cells were counted manually. To determine cell adhesion areas, images (phalloidin stainings of the actin cytoskeleton) were analyzed by thresholding and watershedding. The area of cell clusters was removed when analyzing results. Confocal microscopy images were acquired with a Zeiss 710 Confocal Microscope.

### Human Primary Keratinocyte cell line culture and seeding

Human primary keratinocytes were cultured in Keratinocytes Basal Medium 2 (KBM2, PromoCell). For proliferation assays, HPK cells were harvested with trypsin (0.25 %) and versene solutions (Thermo Fisher Scientific, 0.2 g/L EDTA Na_4_ in Phosphate Buffered Saline) at a ratio of 1/9. Cells were then resuspended with differentiation medium (FAD) prepared with DMEM/F12 (1:1) (1x) and DMEM (Thermofisher Scientific) at a ratio of 1:1, containing 1% L-Glutamine (200 mM), 1% Penicillin-Streptomycin (5,000 U/mL), 0.1% insulin, 0.1% Hydrocortisone equivalent (HCE) and 10% of Fetal Bovine Serum (FBS, Labtech). HPK cells were centrifuged for 5 min at 1200 rpm, counted and resuspended in KBM2, at the desired density before seeding onto substrates. Cells were left to adhere and proliferate in an incubator (37 °C and 5 % CO2) for different time points (at day three and day seven of culture), prior to staining and imaging. For cell spreading assays, HPK cells were harvested and seeded onto fluorinated droplets at a density of 25,000 cells per well (13,000 cell/cm^2^). For passaging, cells were reseeded in a preconditioned T75 flask, with collagen type I (20 μL of collagen into 10 mL of PBS for 20 min), at a density of 250,000 cells per flask.

### Statistical analysis

Statistical analysis was carried out using OriginPro 9 through one-way ANOVA with Tukey test for posthoc analysis. Significance was determined by * P < 0.05, ** P < 0.01, *** P < 0.001 and n.s., non-significant. A full summary of statistical analysis is provided in the supplementary information.

## Supporting Information

Supporting Information is available from the authors.

## Acknowledgements

The authors are grateful for experimental support and training from Dr Dexu Kong. Funding for this work from the European Research Council (ProLiCell, 772462; ProBioFac, 966740) and a Cyprus Scholarship (IKYK, 841C18) is gratefully acknowledged.

## References

[1] H. Tavassoli, S. N. Alhosseini, A. Tay, P. P. Y. Chan, S. K. Weng Oh, M. E. Warkiani, Biomaterials 2018, 181, 333.

[2] F. F. dos Santos, P. Z. Andrade, C. L. da Silva, J. M. S. Cabral, Biotechnol. J. 2013, 8, 644.

[3] L. Peng, J. E. Gautrot, Mater. Today Bio 2021, 12, 100159.

[4] D. Kong, W. Megone, K. D. Q. Nguyen, S. Di Cio, M. Ramstedt, J. E. Gautrot, Nano Lett. 2018, 18, 1946.

[5] D. Kong, L. Peng, S. Di Cio, P. Novak, J. E. Gautrot, ACS Nano 2018, 12, 9206.

[6] D. Kong, L. Peng, M. Bosch-Fortea, A. Chrysanthou, C. V. J.-M. Alexis, C. Matellan, A. Zarbakhsh, G. Mastroianni, A. del Rio Hernandez, J. E. Gautrot, Biomaterials 2022, 284, 121494.

[7] D. Carter, X. He, S. Munson, P. Twigg, K. Gernert, M. Broom, T. Miller, Science 1989, 244, 1195.

[8] B. X. Huang, H.-Y. Kim, C. Dass, J. Am. Soc. Mass Spectrom. 2004, 15, 1237.

[9] M. Fasano, S. Curry, E. Terreno, M. Galliano, G. Fanali, P. Narciso, S. Notari, P. Ascenzi, IUBMB Life Int. Union Biochem. Mol. Biol. Life 2005, 57, 787.

[10] X. M. He, D. C. Carter, Nature 1992, 358, 209.

[11] S. Damodaran, J. Food Sci. 2006, 70, R54.

[12] J. J. Babcock, L. Brancaleon, Int. J. Biol. Macromol. 2013, 53, 42.

[13] D. K. Layman, B. Lönnerdal, J. D. Fernstrom, Nutr. Rev. 2018, 76, 444.

[14] K. L. Jakopović, I. Barukčić, R. Božanić, Mljekarstvo 2016, 66, 3.

[15] A. R. Madureira, C. I. Pereira, A. M. P. Gomes, M. E. Pintado, F. Xavier Malcata, Food Res. Int. 2007, 40, 1197.

[16] X. Yuan, X. Li, X. Zhang, Z. Mu, Z. Gao, L. Jiang, Z. Jiang, J. Food Eng. 2018, 223, 116.

[17] A. Qayum, M. Hussain, M. Li, J. Li, R. Shi, T. Li, A. Anwar, Z. Ahmed, J. Hou, Z. Jiang, Food Hydrocoll. 2021, 110, 106122.

[18] A. F. S. A., Habeeb, Arch. Biochem. Biophys. 1967, 121, 652.

[19] Jamileh. Lakkis, Ricardo. Villota, J. Agric. Food Chem. 1992, 40, 553.

[20] V. Vetri, F. Librizzi, M. Leone, V. Militello, Eur. Biophys. J. 2007, 36, 717.

[21] W. Zhong, M. Wen, J. Xu, H. Wang, L.-L. Tan, L. Shang, Chem. Commun. 2020, 56, 11414.

[22] F. Jiménez-Ángeles, H.-K. Kwon, K. Sadman, T. Wu, K. R. Shull, M. Olvera de la Cruz, ACS Cent. Sci. 2019, 5, 688.

[23] J. A. Jamison, E. L. Bryant, S. B. Kadali, M. S. Wong, V. L. Colvin, K. S. Matthews, M. K. Calabretta, J. Nanoparticle Res. 2011, 13, 625.

[24] C. Ma, A. Malessa, A. J. Boersma, K. Liu, A. Herrmann, Adv. Mater. 2020, 32, 1905309.

[25] A. Korpi, C. Ma, K. Liu Nonappa, A. Herrmann, O. Ikkala, M. A. Kostiainen, ACS Macro Lett. 2018, 7, 318.

[26] E. te Brinke, J. Groen, A. Herrmann, H. A. Heus, G. Rivas, E. Spruijt, W. T. S. Huck, Nat. Nanotechnol. 2018, 13, 849.

[27] K. Liu, D. Pesce, C. Ma, M. Tuchband, M. Shuai, D. Chen, J. Su, Q. Liu, J. Y. Gerasimov, A. Kolbe, W. Zajaczkowski, W. Pisula, K. Müllen, N. A. Clark, A. Herrmann, Adv. Mater. 2015, 27, 2459.

[28] J. Bergfreund, M. Diener, T. Geue, N. Nussbaum, N. Kummer, P. Bertsch, G. Nyström, P. Fischer, Soft Matter 2021, 17, 1692.

[29] M. A. Bos, T. van Vliet, Adv. Colloid Interface Sci. 2001, 91, 437.

[30] J. Benjamins, J. Lyklema, E. H. Lucassen-Reynders, Langmuir 2006, 22, 6181.

[31] R. Feng, Y. Konishi, A. W. Bell, J. Am. Soc. Mass Spectrom. 1991, 2, 387.

[32] I. M. Vlasova, A. M. Saletsky, J. Appl. Spectrosc. 2009, 76, 536.

[33] H. T. M. Phan, S. Bartelt-Hunt, K. B. Rodenhausen, M. Schubert, J. C. Bartz, PLOS ONE 2015, 10, e0141282.

[34] R. Li, Z. Wu, Y. Wangb, L. Ding, Y. Wang, Biotechnol. Rep. 2016, 9, 46.

[35] B. A. Russell, B. Jachimska, I. Kralka, P. A. Mulheran, Y. Chen, J. Mater. Chem. B 2016, 4, 6876.

[36] E. A. Permyakov, L. J. Berliner, FEBS Lett. 2000, 473, 269.

[37] J. Spöttel, J. Brockelt, S. Falke, S. Rohn, Molecules 2021, 26, 6247.

[38] J. Bergfreund, P. Bertsch, S. Kuster, P. Fischer, Langmuir 2018, 34, 4929.

[39] C. J. Pace, J. Gao, Acc. Chem. Res. 2013, 46, 907.

[40] S. Dominguez-Medina, J. Blankenburg, J. Olson, C. F. Landes, S. Link, ACS Sustain. Chem. Eng. 2013, 1, 833.

[41] X. Wang, G. Herting, I. Odnevall Wallinder, E. Blomberg, Phys. Chem. Chem. Phys. 2015, 17, 18524.

[42] J. A. Lori, T. Hanawa, Spectroscopy 2004, 18, 545.

[43] D.-H. Tsai, F. W. DelRio, A. M. Keene, K. M. Tyner, R. I. MacCuspie, T. J. Cho, M. R. Zachariah, V. A. Hackley, Langmuir 2011, 27, 2464.

[44] S. G. Baldursdottir, M. S. Fullerton, S. H. Nielsen, L. Jorgensen, Colloids Surf. B Biointerfaces 2010, 79, 41.

[45] Highberger, John H, Boric Acid Color React. Flavone Deriv. 1939, 61, 2302.

[46] S. Hattori, E. Adachi, T. Ebihara, T. Shirai, I. Someki, S. Irie, J. Biochem. 1999, 125, 9.

[47] J. T. Connelly, J. E. Gautrot, B. Trappmann, D. W.-M. Tan, G. Donati, W. T. S. Huck, F. M. Watt, Nat. Cell Biol. 2010, 12, 711.

[48] B. Trappmann, J. E. Gautrot, J. T. Connelly, D. G. T. Strange, Y. Li, M. L. Oyen, M. A. Cohen Stuart, H. Boehm, B. Li, V. Vogel, J. P. Spatz, F. M. Watt, W. T. S. Huck, Nat. Mater. 2012, 11, 642.

[49] R. A. F. Clark, J. M. Folkvord, R. L. Wertz, J. Invest. Dermatol. 1985, 84, 378.

[50] E. J. O’Keefe, R. E. Payne, N. Russell, D. T. Woodley, J. Invest. Dermatol. 1985, 85, 125.

[51] A. Takashima, F. Grinnell, J. Invest. Dermatol. 1984, 83, 352.

[52] P. H. J. Fiona M. Watt, 1993, 73, 713.

[53] P. Kanchanawong, G. Shtengel, A. M. Pasapera, E. B. Ramko, M. W. Davidson, H. F. Hess, C. M. Waterman, Nature 2010, 468, 580.

[54] P. Pandey, W. Hawkes, J. Hu, W. V. Megone, J. Gautrot, N. Anilkumar, M. Zhang, L. Hirvonen, S. Cox, E. Ehler, J. Hone, M. Sheetz, T. Iskratsch, Dev. Cell 2018, 44, 326.

[55] M. Kim, B. Gweon, U. Koh, Y. Cho, D. W. Shin, M. Noh, J. H. Shin, Biomed. Eng. Lett. 2015, 5, 194.

[56] M. Bennett, M. Cantini, J. Reboud, J. M. Cooper, P. Roca-Cusachs, M. Salmeron-Sanchez, Proc. Natl. Acad. Sci. 2018, 115, 1192.

[57] S. Khetan, M. Guvendiren, W. R. Legant, D. M. Cohen, C. S. Chen, J. A. Burdick, Nat. Mater. 2013, 12, 458.

[58] D. Kong, K. D. Q. Nguyen, W. Megone, L. Peng, J. E. Gautrot, Faraday Discuss. 2017, 204, 367.

[59] L. Pieuchot, J. Marteau, A. Guignandon, T. Dos Santos, I. Brigaud, P.-F. Chauvy, T. Cloatre, A. Ponche, T. Petithory, P. Rougerie, M. Vassaux, J.-L. Milan, N. Tusamda Wakhloo, A. Spangenberg, M. Bigerelle, K. Anselme, Nat. Commun. 2018, 9, 3995.

