## Supplementary Information for "Supercharged Protein Nanosheets for Cell Expansion on Bioemulsions"

Alexandra Chrysanthou[1,2], Hassan Kanso[1,2], Wencheng Zhong[3], Li Shang[3,4] and Julien E. Gautrot[1,2*]

*[1] Institute of Bioengineering and [2]School of Engineering and Materials Science, Queen Mary, University of London, Mile End Road, London, E1 4NS, UK.*

*[3] State Key Laboratory of Solidification Processing, School of Materials Science and Engineering, Northwestern Polytechnical University and Shaanxi Joint Laboratory of Graphene (NPU), Xi'an, 710072, China.*

*[4] NPU-QMUL Joint Research Institute of Advanced Materials and Structures (JRI-AMAS), Northwestern Polytechnical University, Xi'an, 710072, China.*

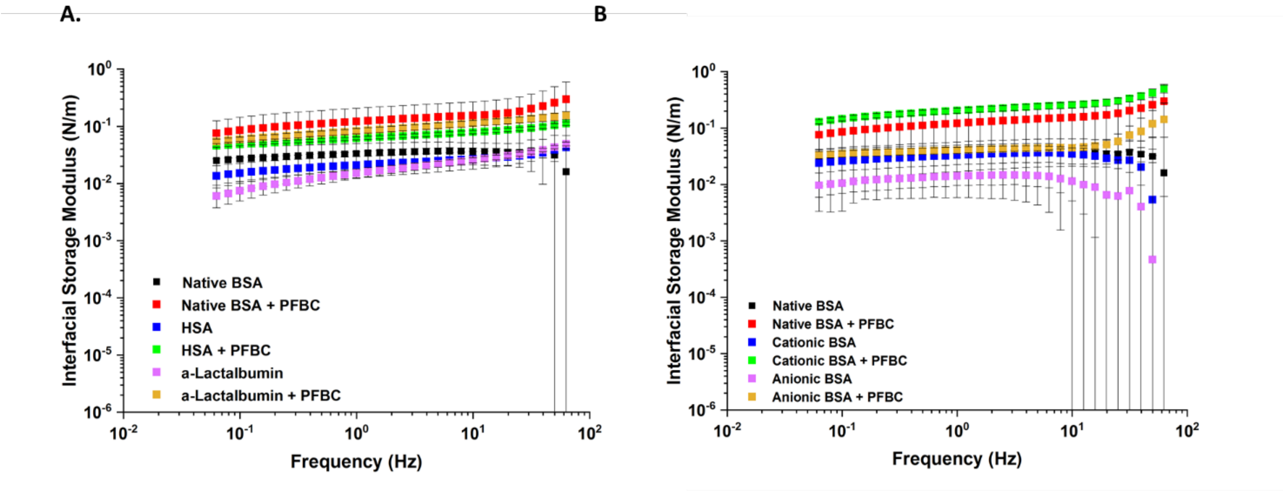

**Supplementary Figure S1.** Frequency sweep at oscillating amplitude of $10^{-4}$ rad carried out for the characterization of (A) BSA, HSA and $\alpha$-Lactalbumin (all at 1 mg/mL) with and without pro-surfactant PFBC and (B) native, cationic (cBSA) and anionic BSA (aBSA) with and without pro-surfactant PFBC (10 $\mu$g/mL). All experiments were carried out at interfaces between PBS and Novec 7500 (fluorinated) oil.

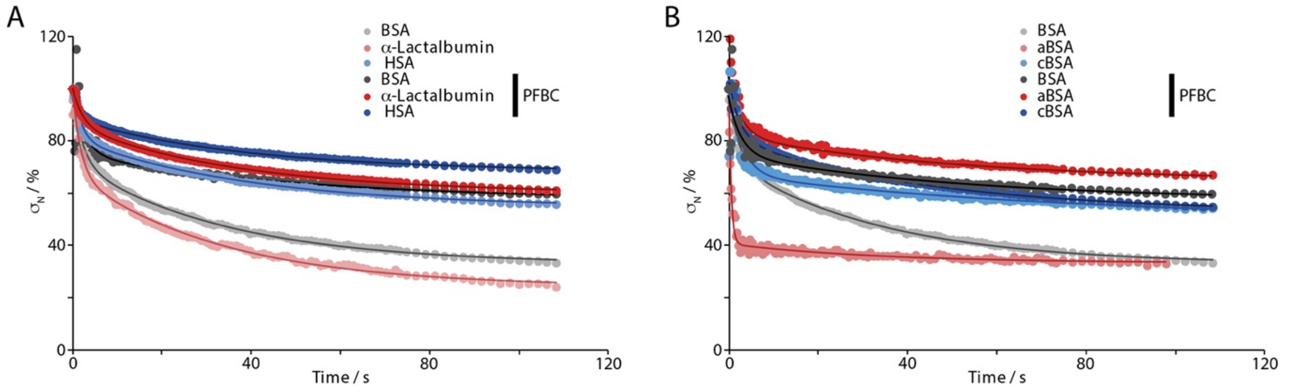

**Supplementary Figure S1.** Normalised stress relaxation experiments carriet out on protein nanosheets formed at liquid-liquid interfaces with and without PFBC (10 $\mu$g/mL). Data is shown as normalised stress ($\sigma_N$) extracted from stress relaxation experiments at a strain 0.5%. (A) Comparison of BSA, HSA, $\alpha$-Lactalbumin and (B) native BSA, cBSA and aBSA.

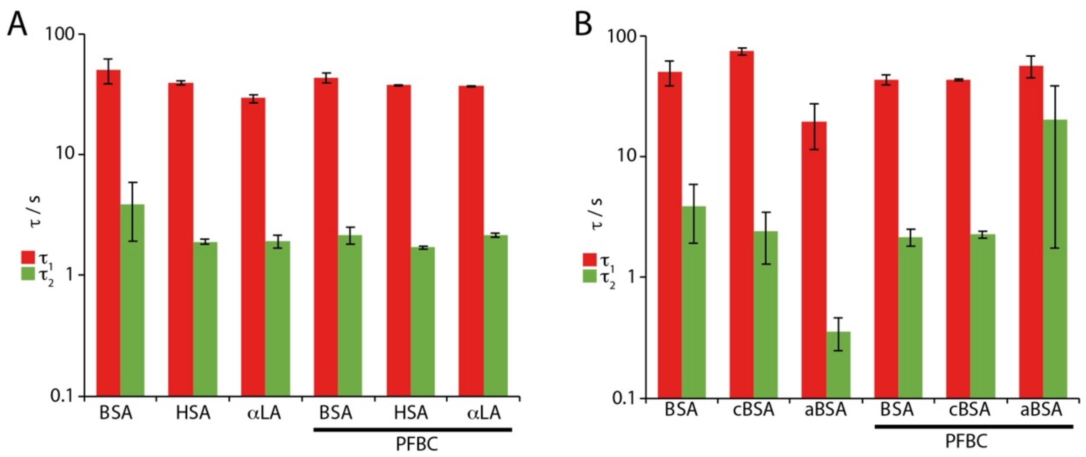

**Supplementary Figure S3.** Characterisation of the stress relaxation profiles associated with protein nanosheets studied. Data were extracted from stress relaxation at a strain of 0.5%. (A) Comparing BSA, HSA, $\alpha$-Lactalbumin (all at 1 mg/mL) and (B) native BSA, cBSA and aBSA. Error bars are s.e.m; n=3.

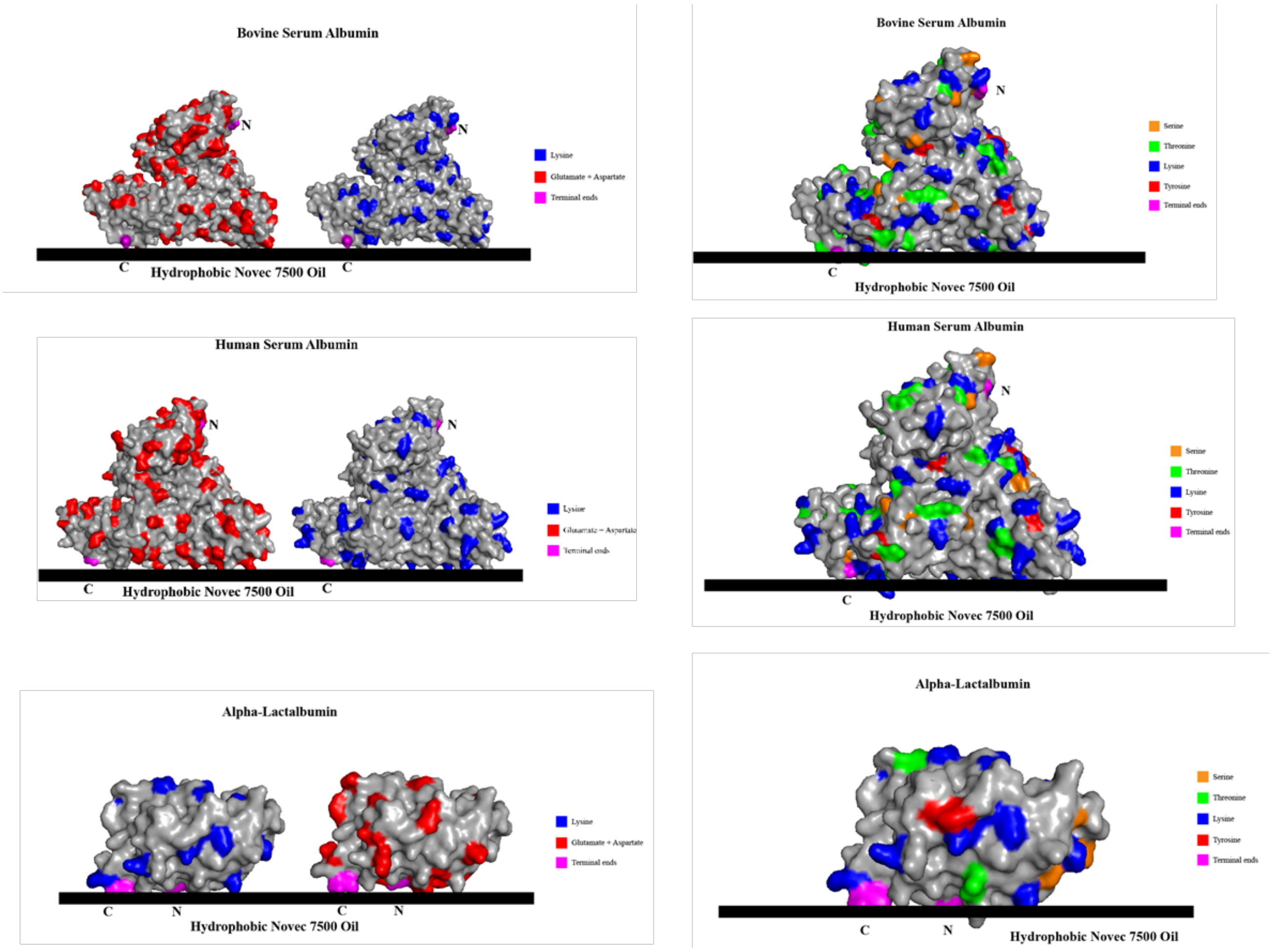

**Supplementary Figure S4.** Proposed model for the adsorption of different albumin proteins at hydrophobic Novec 7500 oil interfaces. Orientations proposed are based on the hydrophobicity of corresponding protein surfaces and are not further optimised via molecular dynamics. Note that it is likely that significant denaturation and remodelling takes place following on from such initial interaction configuation. Left images. Surface accessible residues potentially reacting during the supercharging of corresponding proteins; Red – Glutamate and Aspartate residues, Blue – lysine residues. Pink - Terminal ends. Right images. Surface accessible hydroxyl-based amino acids, and in blue lysine residues. Images were orientated and generated using Pymol software; the draw “Connolly surfaces” command was used to trace the surface of the proteins that are proposed to be in direct contact with the buffer.

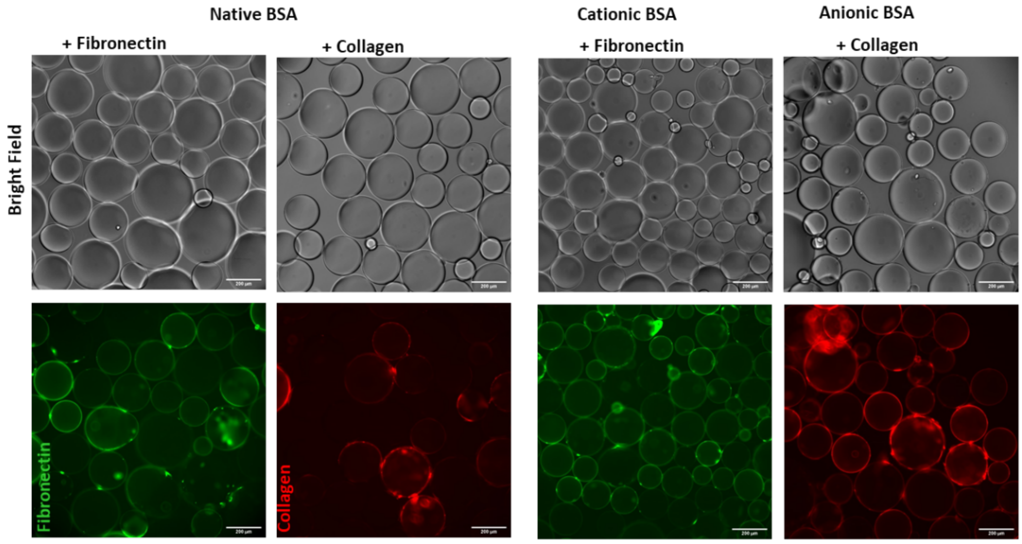

**Supplementary Figure S5.** Epifluorescence microscopy images were with rabbit (green, AB F3648) and mouse (red, AB ab90395) antibodies and species specific secondary antibodies conjugated. Scale bars, 200 μm.

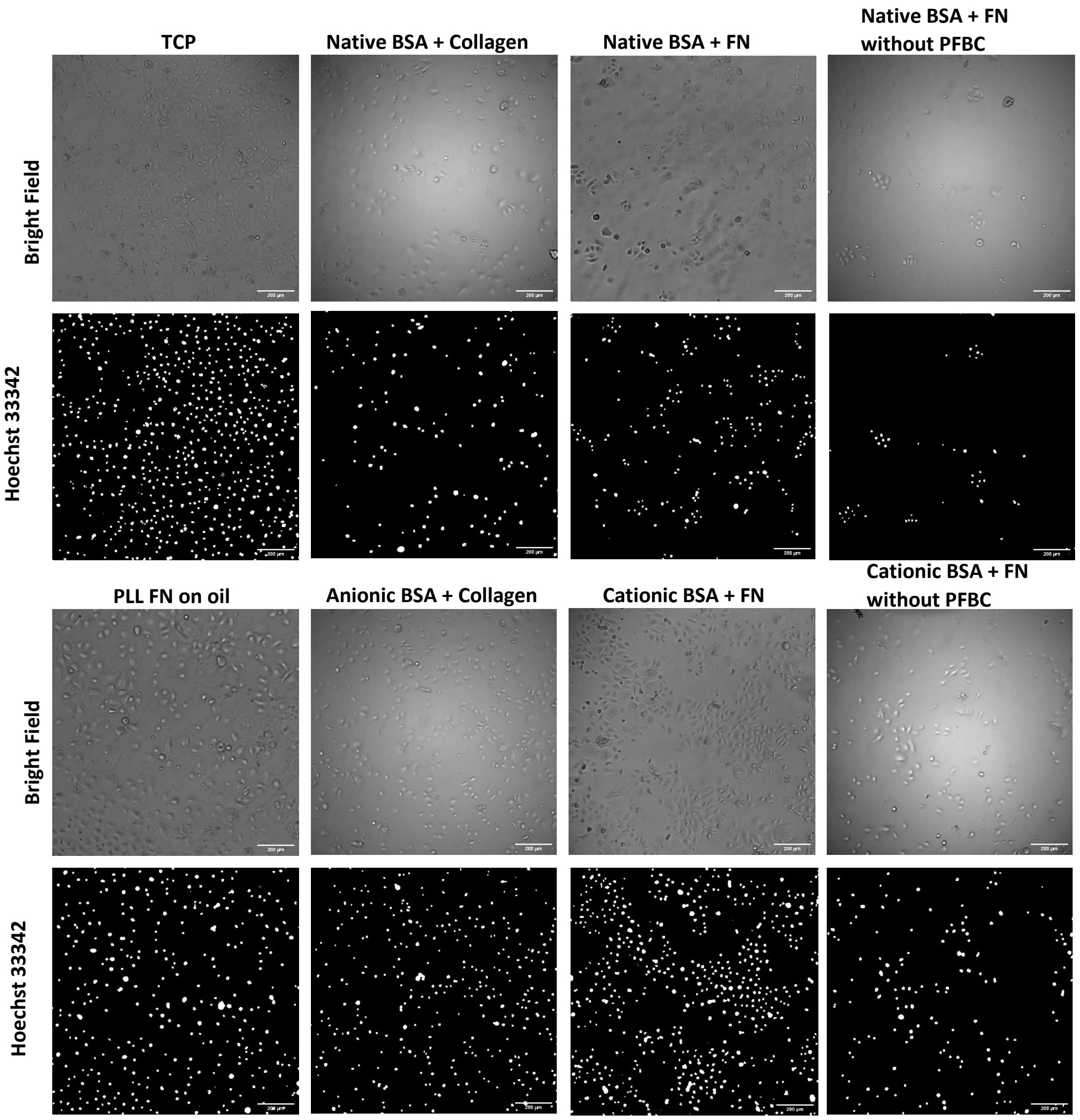

**Supplementary Figure S6.** Bright field and epifluorescence images of HPKs adhered on pinned droplets after three days in culture. Images are corresponding nuclear stainings (Hoechst 33342). Scale bar, 200 µm.

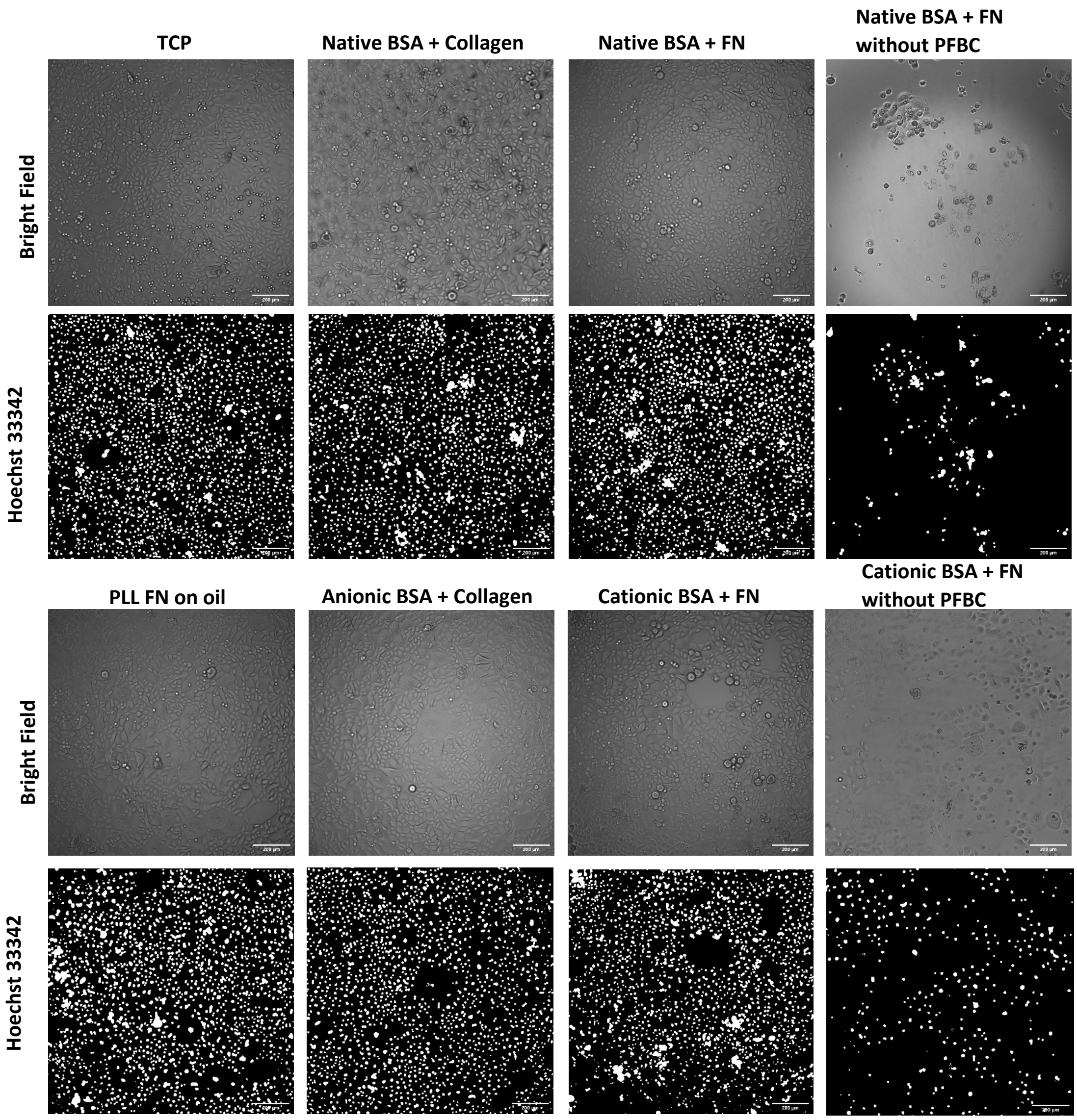

**Supplementary Figure S7.** Bright field and epifluorescence images of HPKs adhered on pinned droplets after seven days in culture. Images are corresponding nuclear stainings (Hoechst 33342). Scale bars, 200 µm.

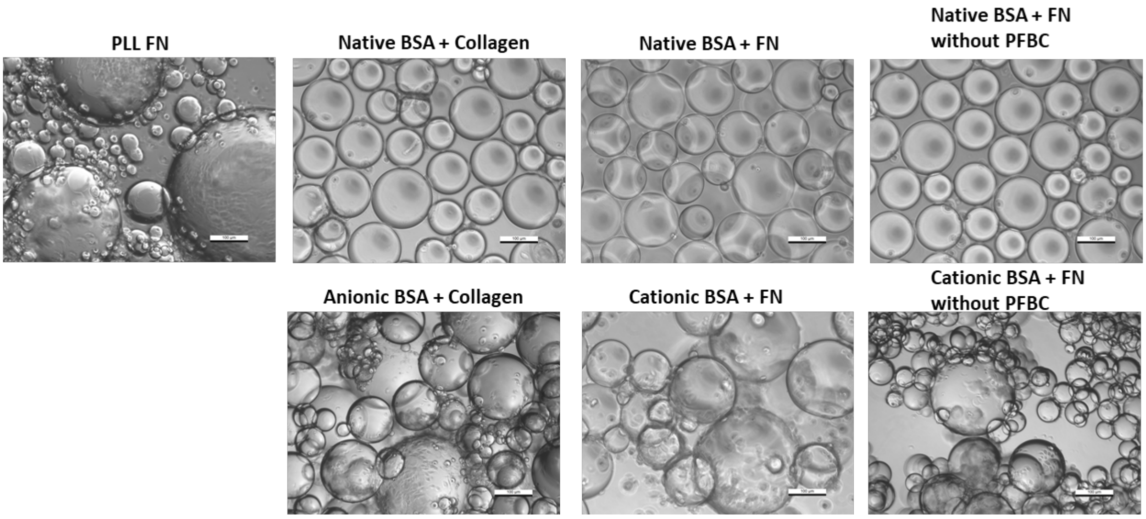

**Supplementary Figure S8.** Bright field images showing the HPKs adhesion and growing on the oil droplets after three days of culture. Scale bars, 100 µm.

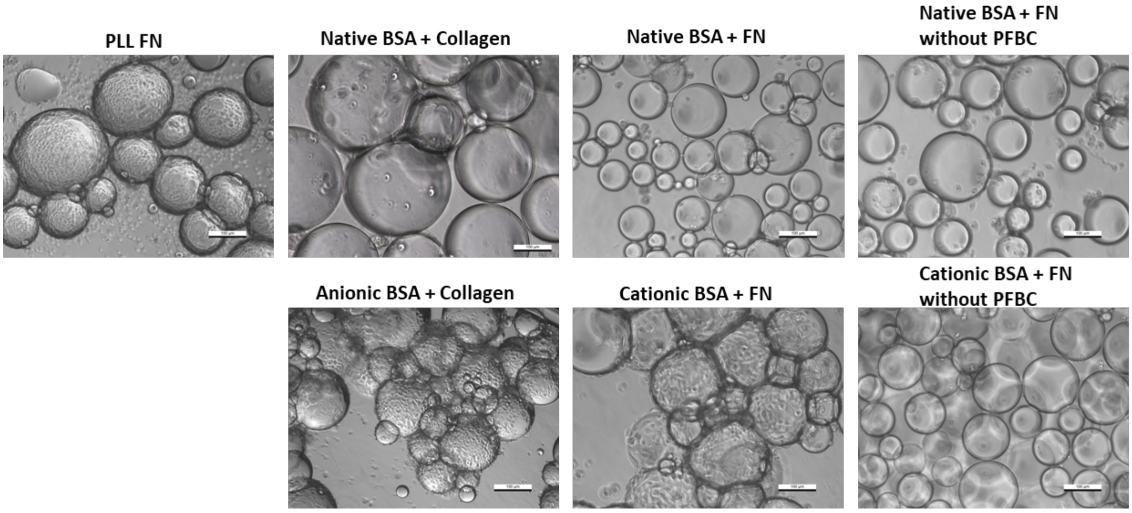

**Supplementary Figure S9.** Bright field images showing the HPKs adhesion and growing on the oil droplets after seven days of culture. Scale bars, 100 µm.

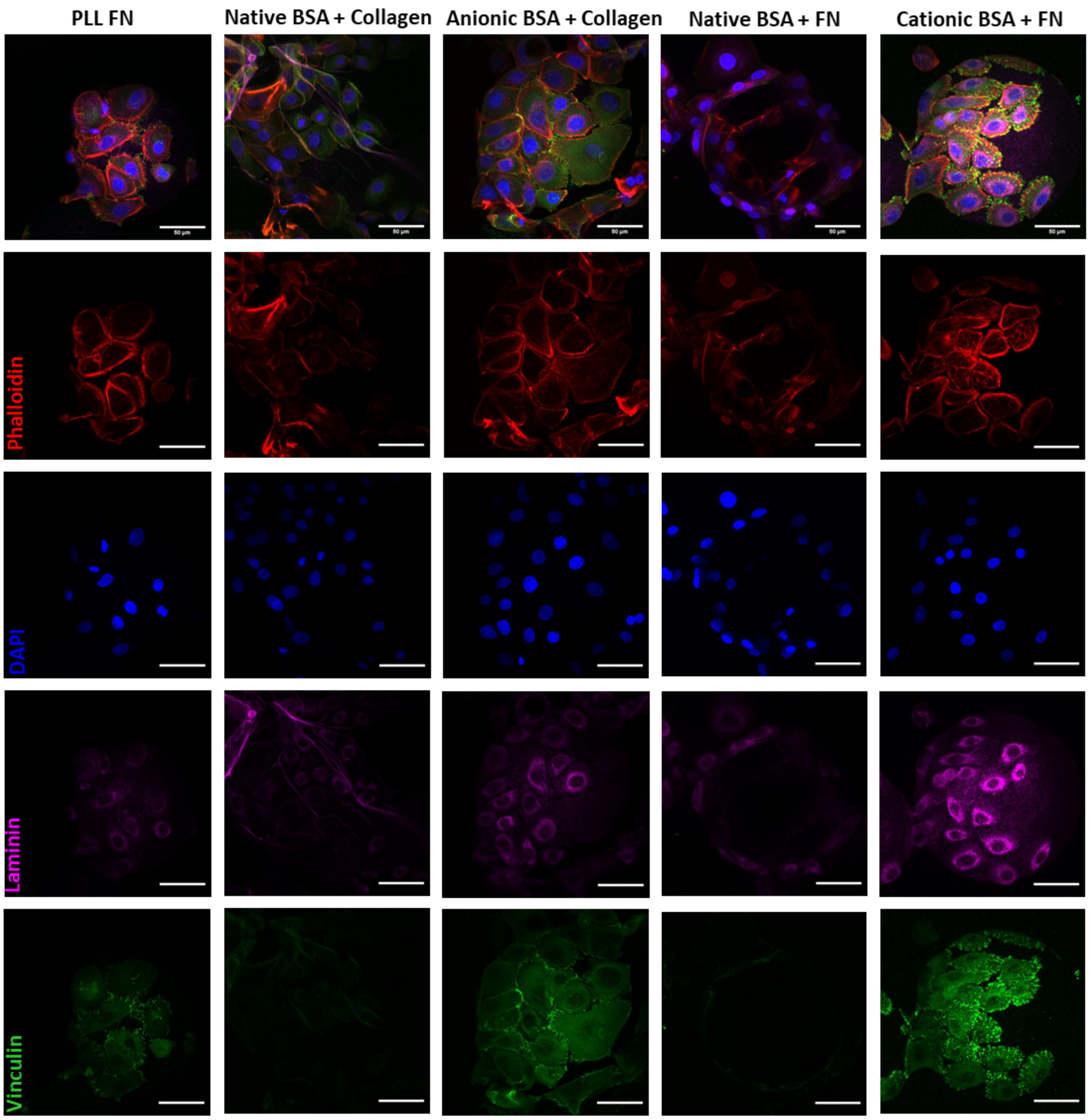

**Supplementary Figure S10.** Confocal fluorescence microscopy images of HPKs spreading after seven days on emulsion (blue, DAPI; red, phalloidin; green, vinculin; purple, laminin). Error bars are s.e.m.; n = 3. Scale bars, 50 µm.

### Supplementary Tables

**Supplementary Table S1.** Summary of statistical analysis of data obtained by frequency sweep after the proteins were adsorbed at fluorinated-PBS interfaces with and without co-surfactant (PFBC) at 10 μg/ mL (Figures 1E and 2E).

| | **MeanDiff** | **Prob** | |
|---|---|---|---|
| Cationic BSA Native BSA | -0.02877 | 1.58E-04 | *** |
| Anionic BSA Native BSA | -0.04202 | 5.37E-06 | *** |
| Anionic BSA Cationic BSA | -0.01325 | 0.03919 | * |
| α-Lactalbumin Native BSA | -0.01816 | 0.00563 | ** |
| α-Lactalbumin Cationic BSA | 0.01061 | 0.11291 | n.s |
| α-Lactalbumin Anionic BSA | 0.02385 | 7.38E-04 | *** |
| HSA Native BSA | -0.01692 | 0.00907 | ** |
| HSA Cationic BSA | 0.01185 | 0.06883 | n.s |
| HSA Anionic BSA | 0.0251 | 4.91E-04 | *** |
| HSA α-Lactalbumin | 0.00124 | 0.99721 | n.s |
| Cationic BSA with PFBC Native BSA with PFBC | 0.21018 | 3.49E-07 | *** |
| Anionic BSA with PFBC Native BSA with PFBC | -4.25E-04 | 1 | n.s |
| Anionic BSA with PFBC Cationic BSA with PFBC | -0.21061 | 3.43E-07 | *** |
| α-Lactalbumin with PFBC Native BSA with PFBC | 0.06796 | 0.0055 | ** |
| α-Lactalbumin with PFBC Cationic BSA with PFBC | -0.14222 | 1.31E-05 | *** |
| α-Lactalbumin with PFBC Anionic BSA with PFBC | 0.06839 | 0.00527 | ** |
| HSA with PFBC Native BSA with PFBC | 0.03371 | 0.20575 | n.s |
| HSA with PFBC Cationic BSA with PFBC | -0.17648 | 2.23E-06 | *** |
| HSA with PFBC Anionic BSA with PFBC | 0.03413 | 0.19724 | n.s |
| HSA with PFBC α-Lactalbumin with PFBC | -0.03425 | 0.19491 | n.s |

**Supplementary Table S2.** Summary of statistical analysis of data obtained by frequency sweep after the proteins were adsorbed at the fluorinated-PBS interface with and without the surfactant at 10 μg/ mL. Pairwise comparison to determine the significance of the effect of prosurfactant (Figures 1E and 2E).

| | **MeanDiff** | **Prob** | |
|---|---|---|---|
| Native BSA with PFBC Native BSA | 0.00165 | 0.57633 | n.s |
| Cationic BSA with PFBC Cationic BSA | 0.24061 | 3.42E-04 | *** |
| Anionic BSA with PFBC Anionic BSA | 0.04324 | 2.39E-07 | *** |
| α-Lactalbumin with PFBC α-Lactalbumin | 0.08778 | 3.15E-04 | *** |
| HSA with PFBC HSA | 0.05228 | 9.98E-04 | *** |

**Supplementary Table S3.** Summary of statistical analysis of data obtained from stress relaxation experiments at a strain of 0.5 % (Figures 1F and 2F).

| | MeanDiff | Prob | |
|---|---|---|---|
| BSA PFBC BSA | 24.309 | 0.03934 | * |
| cBSA PFBC cBSA | 4.34233 | 0.63335 | n.s |
| aBSA PFBC aBSA | 36.47233 | 0.02059 | * |
| HSA PFBC HSA | 16.679 | 0.05404 | n.s |
| $\alpha$-Lactalbumin PFBC $\alpha$-Lactalbumin | 41.525 | 0.0022 | ** |

**Supplementary Table S4.** Summary of statistical analysis of data obtained from the SPR data for the protein binding (1 mg/mL) at the perfluorodecanethiol pre-treated chips (Figure 3B).

| | MeanDiff | Prob | |
|---|---|---|---|
| Anionic BSA Native BSA | 296.9 | 0.07194 | n.s |
| Cationic BSA Native BSA | 1381.267 | 3.27E-05 | *** |
| Cationic BSA Anionic BSA | 1084.367 | 1.32E-04 | *** |

**Supplementary Table S5.** Summary of statistical analysis of data obtained from the SPR data for the fibronectin or collagen binding at the surface of supercharged protein layers (native, cationic and anionic BSA) (Figure 3D).

| | MeanDiff | Prob | |
|---|---|---|---|
| BSA/Col BSA/FN | -37.3333 | 0.98415 | n.s |
| cBSA/FN BSA/FN | -30 | 0.99159 | n.s |
| cBSA/FN BSA/Col | 7.33333 | 0.99987 | n.s |
| aBSA/Col BSA/FN | 679 | 9.71E-04 | *** |
| aBSA/Col BSA/Col | 716.3333 | 6.76E-04 | *** |
| aBSA/Col cBSA/FN | 709 | 7.25E-04 | *** |

**Supplementary Table S6.** Summary of statistical analysis of data obtained from the epifluorescence images for the fibronectin or collagen binding on native, anionic and cationic BSA emulsion droplets (Figure 3F).

| | MeanDiff | Prob | |
|---|---|---|---|
| Cationic BSA + FN Native BSA + FN | 4425.399 | 0.00282 | ** |
| Anionic BSA + Collagen Native BSA + Collagen | 6989.911 | 0.00253 | ** |
| Native BSA + Collagen Native BSA + FN | -2893.23 | 0.02859 | * |
| Anionic BSA + Collagen Cationic BSA + FN | -328.722 | 0.72984 | n.s |

**Supplementary Table S7.** Summary of statistical analysis of data obtained from cell (HPK) density on pinned droplets after three days in culture (Figure 4A).

| | MeanDiff | Prob | |
|---|---|---|---|
| PLL FN TPS | -104.667 | 0.90088 | n.s |
| Native BSA Collagen TPS | -432.444 | 0.00173 | * |
| Native BSA Collagen PLL FN | -327.778 | 0.0199 | * |
| Anionic BSA Collagen TPS | -351.667 | 0.01138 | * |
| Anionic BSA Collagen PLL FN | -247 | 0.12135 | n.s |
| Anionic BSA Collagen Native BSA Collagen | 80.77778 | 0.97227 | n.s |
| Native BSA FN TPS | -354.556 | 0.01064 | * |
| Native BSA FN PLL FN | -249.889 | 0.11422 | n.s |
| Native BSA FN Native BSA Collagen | 77.88889 | 0.9772 | n.s |
| Native BSA FN Anionic BSA Collagen | -2.88889 | 1 | n.s |
| Cationic BSA FN TPS | -201.111 | 0.29522 | n.s |
| Cationic BSA FN PLL FN | -96.4444 | 0.93205 | n.s |
| Cationic BSA FN Native BSA Collagen | 231.3333 | 0.16716 | n.s |
| Cationic BSA FN Anionic BSA Collagen | 150.5556 | 0.62167 | n.s |
| Cationic BSA FN Native BSA FN | 153.4444 | 0.601 | n.s |
| Native BSA FN no PFBC TPS | -534.111 | 1.78E-04 | *** |
| Native BSA FN no PFBC PLL FN | -429.444 | 0.00185 | ** |
| Native BSA FN no PFBC Native BSA Collagen | -101.667 | 0.91308 | n.s |
| Native BSA FN no PFBC Anionic BSA Collagen | -182.444 | 0.40249 | n.s |
| Native BSA FN no PFBC Native BSA FN | -179.556 | 0.42085 | n.s |
| Native BSA FN no PFBC Cationic BSA FN | -333 | 0.01762 | * |
| Cationic BSA FN no PFBC TPS | -460 | 9.18E-04 | *** |
| Cationic BSA FN no PFBC PLL FN | -355.333 | 0.01044 | * |
| Cationic BSA FN no PFBC Native BSA Collagen | -27.5554 | 0.99997 | n.s |
| Cationic BSA FN no PFBC Anionic BSA Collagen | -108.333 | 0.88469 | n.s |
| Cationic BSA FN no PFBC Native BSA FN | -105.444 | 0.89756 | n.s |
| Cationic BSA FN no PFBC Cationic BSA FN | -258.889 | 0.09432 | n.s |
| Cationic BSA FN no PFBC Native BSA FN no PFBC | 74.11122 | 0.98265 | n.s |

**Supplementary Table S8.** Summary of statistical analysis of data obtained from cell (HPK) density on pinned droplets after seven days in culture (Figure 4A).

| | MeanDiff | Prob | |
|---|---|---|---|
| PLL FN TPS | -129.333 | 0.99923 | n.s |
| Native BSA Collagen TPS | -461.778 | 0.56631 | n.s |
| Native BSA Collagen PLL FN | -332.444 | 0.85717 | n.s |
| Anionic BSA Collagen TPS | -543.667 | 0.37824 | n.s |
| Anionic BSA Collagen PLL FN | -414.333 | 0.68207 | n.s |
| Anionic BSA Collagen Native BSA Collagen | -81.8889 | 0.99996 | n.s |
| Native BSA FN TPS | -592.889 | 0.28432 | n.s |
| Native BSA FN PLL FN | -463.556 | 0.56197 | n.s |
| Native BSA FN Native BSA Collagen | -131.111 | 0.99916 | n.s |
| Native BSA FN Anionic BSA Collagen | -49.2222 | 1 | n.s |
| Cationic BSA FN TPS | -473.111 | 0.53874 | n.s |
| Cationic BSA FN PLL FN | -343.778 | 0.83641 | n.s |
| Cationic BSA FN Native BSA Collagen | -11.3333 | 1 | n.s |
| Cationic BSA FN Anionic BSA Collagen | 70.55556 | 0.99999 | n.s |
| Cationic BSA FN Native BSA FN | 119.7778 | 0.99953 | n.s |
| Native BSA FN no PFBC TPS | -2030.89 | 6.72E-06 | *** |
| Native BSA FN no PFBC PLL FN | -1901.56 | 1.58E-05 | *** |
| Native BSA FN no PFBC Native BSA Collagen | -1569.11 | 1.64E-04 | *** |
| Native BSA FN no PFBC Anionic BSA Collagen | -1487.22 | 3.01E-04 | *** |
| Native BSA FN no PFBC Native BSA FN | -1438 | 4.38E-04 | *** |
| Native BSA FN no PFBC Cationic BSA FN | -1557.78 | 1.78E-04 | *** |
| Cationic BSA FN no PFBC TPS | -1660.33 | 8.43E-05 | *** |
| Cationic BSA FN no PFBC PLL FN | -1531 | 2.17E-04 | *** |
| Cationic BSA FN no PFBC Native BSA Collagen | -1198.56 | 0.00284 | ** |
| Cationic BSA FN no PFBC Anionic BSA Collagen | -1116.67 | 0.00548 | ** |
| Cationic BSA FN no PFBC Native BSA FN | -1067.44 | 0.00815 | ** |
| Cationic BSA FN no PFBC Cationic BSA FN | -1187.22 | 0.00311 | ** |
| Cationic BSA FN no PFBC Native BSA FN no PFBC | 370.5556 | 0.78223 | n.s |

**Supplementary Table S9.** Summary of statistical analysis of data obtained from cell adhesion area on pinned droplets after 48h in culture (Figure 4C).

| | MeanDiff | Prob | |
|---|---|---|---|
| PLL FN TCP | 238.244 | 0.51286 | n.s |
| Anionic BSA Collagen Native BSA Collagen | 463.9954 | 0.51532 | n.s |
| Cationic BSA FN Native BSA FN | 449.8663 | 0.34017 | n.s |
| Native BSA Collagen PLL FN | -1006.69 | 0.03179 | * |
| Anionic BSA Collagen PLL FN | -542.691 | 0.44682 | n.s |
| Native BSA FN PLL FN | -683.948 | 0.0558 | n.s |
| Cationic BSA FN PLL FN | -234.082 | 0.62417 | n.s |

**Supplementary Table S10.** Summary of statistical analysis of data obtained by mean fluorescence intensity for laminin deposition from cells cultured on emulsion droplets (Figure 4D).

| | MeanDiff | Prob | |
|---|---|---|---|
| Native BSA + Collagen PLL + FN | 0.89844 | 0.91187 | n.s |
| Anionic BSA + Collagen PLL + FN | 7.97922 | 1.63E-04 | *** |
| Anionic BSA + Collagen Native BSA + Collagen | 7.08078 | 4.40E-04 | *** |
| Native BSA + FN PLL + FN | -1.66067 | 0.55494 | n.s |
| Native BSA + FN Native BSA + Collagen | -2.55911 | 0.19424 | n.s |
| Native BSA + FN Anionic BSA + Collagen | -9.63989 | 3.12E-05 | *** |
| Cationic BSA + FN PLL + FN | 12.44644 | 2.99E-06 | *** |
| Cationic BSA + FN Native BSA + Collagen | 11.548 | 6.04E-06 | *** |
| Cationic BSA + FN Anionic BSA + Collagen | 4.46722 | 0.01276 | * |
| Cationic BSA + FN Native BSA + FN | 14.10711 | 1.42E-06 | *** |
